# One-step generation of a conditional allele in mice using a short artificial intron

**DOI:** 10.1101/2022.09.06.506831

**Authors:** Annelise M Cassidy, Destinée B Thomas, Emin Kuliyev, Hanying Chen, Stephane Pelletier

## Abstract

Despite tremendous advances in genome editing technologies, generation of conditional alleles in mice has remained challenging. Recent studies in cells have successfully made use of short artificial introns to engineer conditional alleles. The approach consists of inserting intronic sequences flanked by two loxP sites within an exon of a gene using CRISPR-Cas9 technology. Under normal conditions, the artificial intron is removed by the splicing machinery, allowing for expression of the gene product. Following Cre-mediated recombination of the two loxP sites, the intron is disabled, and splicing can no longer occur. The remaining intronic sequences create a frameshift and early translational termination. Here we describe the application of this technology to engineer conditional alleles in mice using *Scyl1* as a model gene. Insertion of the cassette occurred in 17% of edited mice obtained from pronuclear stage zygote microinjection. Mice homozygous for the insertion expressed SCYL1 at levels comparable to wild-type mice and showed no overt abnormalities associated with the loss of *Scyl1* function, indicating the proper removal of the artificial intron. Deletion of the cassette via Cre-mediated recombination in vivo occurred at high frequency, abrogated SCYL1 protein expression, and resulted in loss-of-function phenotypes. Our results broaden the applicability of this approach to engineering conditional alleles in mice.

## Introduction

Genetically modified mouse models have been instrumental in understanding gene function and modeling human disease. The engineering of mouse models has been greatly simplified with the discovery and implementation of targetable nucleases such as Clustered Regularly Interspaced Short Palindromic Repeats (CRISPR)-CRISPR-associated protein 9 (Cas9) systems for genome editing. These enzymes function by introducing DNA double strand breaks (DSBs) at a precise location within a genome of interest which in turn activate DNA repair pathways that can be overtaken to introduce specific mutations by the coadministration of DNA repair templates (Pelletier et al., 2015).

Despite these technological advances, the generation of mouse models with conditional alleles has remained challenging. Two main strategies are currently used to engineer these models. The first strategy consists of flanking one or more critical exons of a gene with site specific recombinase sequences (SSRSs) such that upon recombination between the two SSRSs, the intervening region is eliminated. Splicing between the remaining exons creates a frame shift and premature a stop codon, leading to early translational termination and degradation of the defective mRNA transcript via the nonsense-mediated mRNA decay pathway (Popp and Maquat, 2016). The second strategy involves the insertion of long and complex artificial introns within an exon of a gene (Andersson-Rolf et al., 2017; Economides et al., 2013). These artificial introns contain sequences encoding head-to-head SSRSs, reporter cassettes, and resistance genes running in the opposite direction of the target gene. Upon recombination between the two SSRSs, the cassette is inverted, and a cryptic splice acceptor is exposed, resulting in both the expression of a reporter gene and inactivation of the target gene. Both strategies can be used for engineering mouse models via conventional or CRISPR-assisted gene targeting in embryonic stem (ES) cells or via microinjection of CRISPR-Cas9 components. While these approaches have been used to engineer thousands of mouse models with conditional alleles, they are inefficient, labor intensive, and very expensive. Moreover, the presence of strong promoters within reporter cassettes may have an impact on expression of surrounding genes (Soulez et al., 2019).

A variation of the artificial intron approach called DECAI (DEgradation based on Cre-regulated-Artificial Intron) was recently described in cultured cells (Guzzardo et al., 2017). In this system, a short DNA cassette containing intronic sequences flanked by two loxP sites is inserted in an exon of a gene. In the absence of Cre recombinase, the artificial intron is removed by the splicing machinery allowing for normal expression of the protein. In the presence of Cre recombinase, the intron is crippled and sequences remain, resulting in premature translational termination and degradation of the faulty mRNA transcript through the nonsense-mediated mRNA decay pathway. While the approach was successful in cultured cells, it has never been used to engineer mouse models. Implementing this strategy may have a significant impact on improving the speed and cost of generating these models. Insertion of a short DNA segment in lieu of large DNA constructs or pairs of loxP sites flanked by homology arms may reduce the number of rounds of microinjection currently required for the generation of these mouse models. Short DNA constructs are also more affordable to produce and their insertion within the genome is easier to characterize.

Here we show the successful application of this approach for engineering conditional alleles in mice. Using *Scyl1* as a prototypic gene, we inserted a small artificial intron within exon 3. The insertion occurred in 16.7% of edited mice obtained from microinjections. Mice homozygous for the insertion expressed SCYL1 to levels that are comparable to wild-type animals, illustrating the functionality of the allele. Incapacitation of the artificial intron by crossing these mice to Cre-deleter mice resulted in the complete inactivation of the allele and induced phenotypic changes consistent with the loss of *Scyl1* function. Mechanistically, we found gene inactivation occurs at both the transcriptional and translational levels.

## Results

In search of more efficient and effective strategies to engineer conditional alleles in mice, we turned to the recently developed DECAI system (Guzzardo *et al*., 2017). This approach consists of inserting a short 201-nucleotide-long DNA cassette within an exon of a gene. The cassette contains sequences encoding a splice donor, stop codons in all three frames, two loxP sites flanking an intronic branch point and a polypyrimidine tract, followed by a splice acceptor (Fig. 1). In the absence of Cre recombinase, the intron is functional and removed by the splicing machinery, allowing for normal expression of the protein. In the presence of Cre recombinase, recombination between the two loxP sites occurs resulting in the removal of the branch point and the polypyrimidine tract, thereby inactivating the intron. The crippled intron is then no longer recognized by the splicing machinery and remains in the transcript, resulting in early translational termination and degradation of the damaged mRNA via the nonsense-mediated mRNA decay pathway.

**Figure 1.**
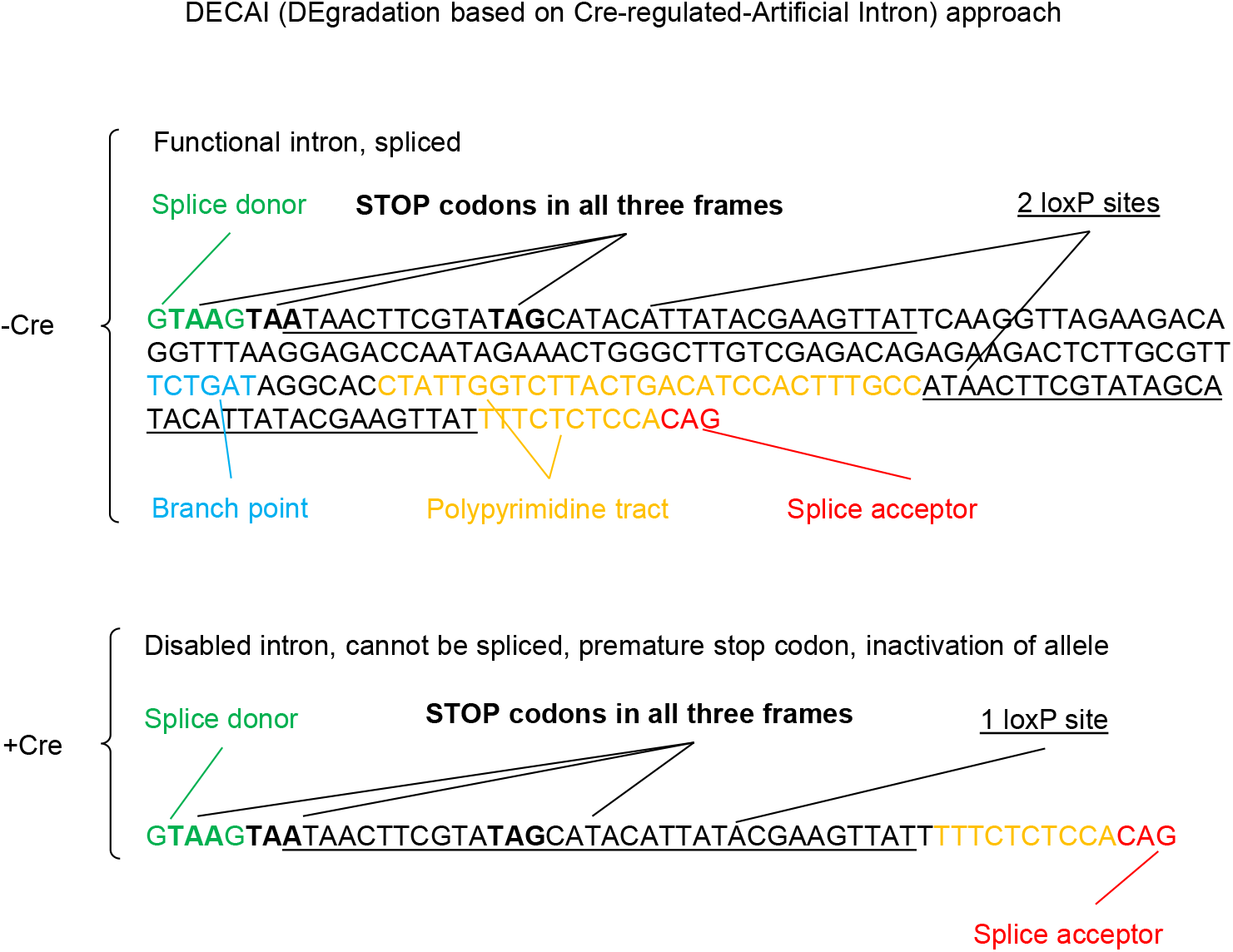
Sequence of the artificial intron used to engineer the *Scyl1^AIv4^* conditional allele. A) The sequence for the artificial intron was derived from (Guzzardo *et al*., 2017), and contains sequences encoding a splice donor (green), followed by stop codons in all three frames (bold), a loxP site (underlined), a branch point (blue), a polypyrimidine tract (yellow), a second loxP site (underlined), and a splice acceptor (red). In the absence of Cre recombinase, the artificial intron is removed by the splicing machinery, allowing translation of the protein. Upon Cre-mediated recombination of the two loxP sites, the artificial intron is disabled such that after transcription and maturation of the transcript, the crippled intron cannot be removed, leading to early termination of translation and degradation of the mRNA transcript via the nonsense-mediated mRNA decay pathway.

To test whether this approach could be effectively used to engineer conditional alleles in mice, the cassette was inserted within exon 3 of the *Scyl1* gene via microinjection of pronuclear stage zygotes with Cas9 mRNA transcripts, a single guide RNA (sgRNA), and a homology-directed repair (HDR) template in the form of a short single stranded DNA (ssDNA) molecule with sequences encoding the artificial intron flanked by homology arms of 58 and 60 nucleotides (Fig. 2A). From two rounds of microinjection, a total of 27 mice were obtained, 12 of which were edited by Cas9, and 2 of which contained the insertion (16.7% of edited mice). Sanger sequencing of TOPO cloned PCR fragments confirmed the proper insertion of the artificial intron (Fig. 2B).

**Figure 2.**
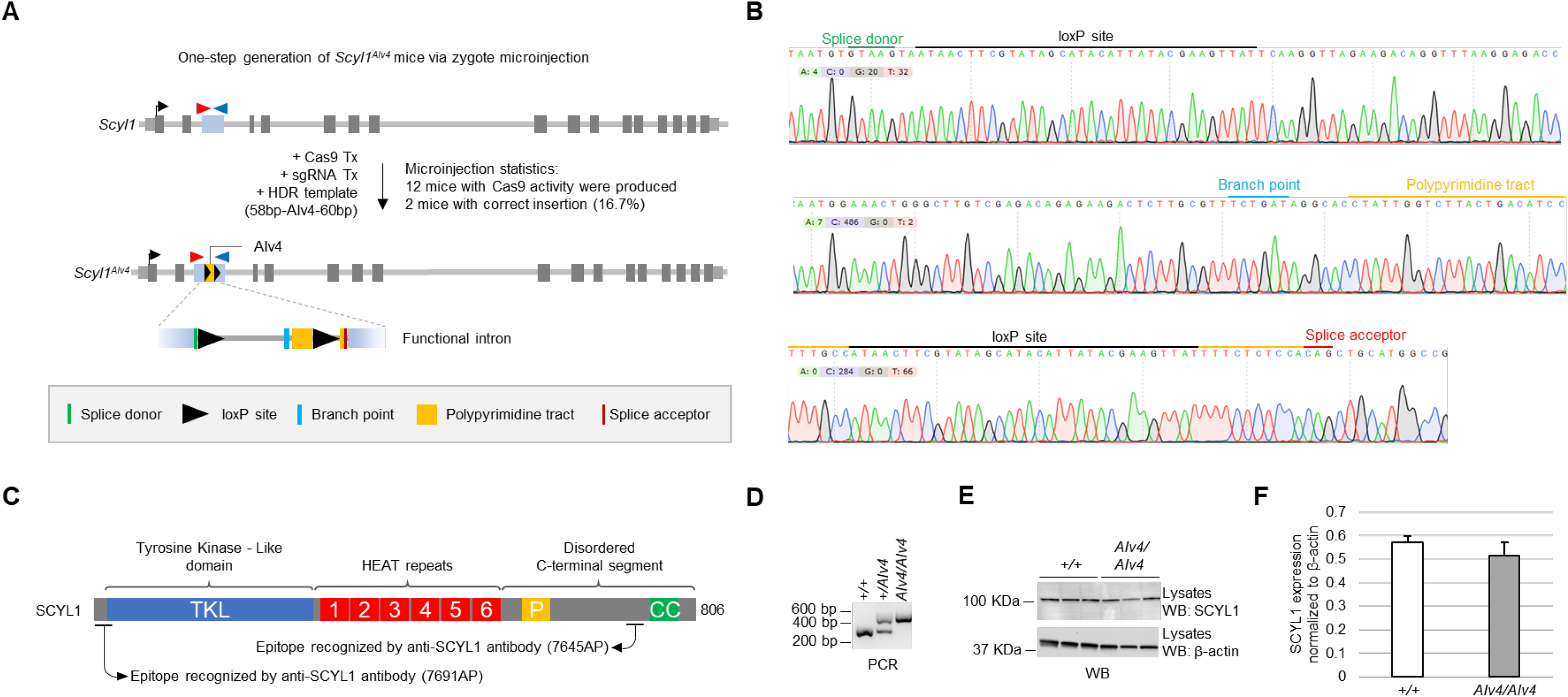
Generation of a *Scyl1* conditional allele in mice using the DECAI approach. A) Schematic representation of the wild type *Scyl1* and *Scyl1^AIv4^* alleles and generation of *Scyl1^AIv4^* mice. Dark gray boxes represent exons, with exon 3 highlighted in blue. Light gray boxes represent the 5’ and 3’ untranslated regions of the gene. Red and blue triangles represent genotyping primers F51 and R52, respectively. The black arrowheads represent the start sites. The yellow box flanked by two black triangles represents the artificial intron inserted into exon 3 of Scyl1. *Scyl1^AIv4^* mice were engineered via pronuclear stage zygote microinjection with Cas9 mRNA transcript, a sgRNA targeting exon 3 of *Scyl1*,and an HDR template containing sequences encoding the artificial intron flanked by homology arms of 58 and 60 nucleotides. 12 mice with Cas9 activity were produced, 2 of which contained the artificial intron (16.7%). B) The sequence composition of the AIv4 allele was confirmed by TOPO cloning and Sanger sequencing. C) Schematic representation of the SCYL1 protein, highlighting the regions recognized by the anti-SCYL1 antibodies. D) PCR-based genotyping of *Scyl1^+/+^, Scyl1^+/AIv4^*, and *Scyl1^AIv4/AIv4^* mice. Amplicons of 304 bp correspond to the wild-type allele, whereas amplicons of 505 bp correspond to the AIv4 allele. E) SCYL1 expression in the brain of *Scyl1*^+/+^ and *Scyl1^AIv4/AIv4^* mice, as detected by western blotting (top panel). β-actin western blot (bottom panel) serves as loading control. F) Quantification of SCYL1 expression in the brain of *Scyl1*^+/+^ and *Scyl1^AIv4/AIv4^* mice, normalized to β-actin. Data is expressed as mean ± SEM.

To test the functionality of the allele, founder mice carrying the insertion were outbred twice to C57BL6/J mice and heterozygous mice produced from these outbreeds were used to generate wild-type (*Scyl1*^+/+^), heterozygous (*Scyl1*^+/*AIv4*^), and homozygous (*Scyl1^AIv4/AIv4^*) mice. These mice were then analyzed at the genomic and protein levels by PCR amplification, Sanger sequencing, and western blotting using an antibody against SCYL1 that recognizes an epitope located within the carboxyl-terminal segment of SCYL1 (Fig. 2C, 2D, 2E, 2F). Fragments of 304 base pairs (bp) corresponding to the wild-type allele were present in *Scyl1*^+/+^ and *Scyl1*^+/*AIv4*^ mice whereas fragments of 505 bp, corresponding to the allele containing the artificial intron, were observed in *Scyl1*^+/*AIv4*^ and *Scyl1^AIv4/AIv4^* mice (Fig. 2D). Sanger sequencing of PCR products obtained from both *Scyl1*^+/+^ and *Scyl1^AIv4/AIv4^* mice confirmed the nature of the alleles (not shown). Importantly, SCYL1 expression was comparable in both *Scyl1*^+/+^ and *Scyl1^AIv4/AIv4^* mice, further supporting the view that the artificial intron is efficiently removed by the splicing machinery allowing for the proper expression of the protein (Fig. 2E and 2F).

To test whether recombination between the two loxP sites would occur despite their proximity to one another (113 bp), *Scyl1^AIv4/AIv4^* mice were crossed with Cre-deleter mice which express Cre recombinase in virtually all tissues (B6.C-Tg(CMV-cre)1Cgn/J, abbreviated CMV-Cre+). From this cross, 33 mice were obtained, 16 of which expressed Cre and harbored a copy of the recombined AIv4 allele (AIv4Δ) (Fig. 3A and 3B). Importantly, a recombination frequency greater than 80% was observed in Cre expressing mice (Fig. 3C). In comparison, the recombination frequency between 2 loxP sites flanking exons 2 to 17 of the *Scyl1* gene (~12.5 kb apart) was less than 50% (Fig. 3A, 3D, and 3E). These findings indicated that the recombination frequency between loxP sites located within 113 bp from each other occurred more efficiently than that of distally located loxP sites.

**Figure 3.**
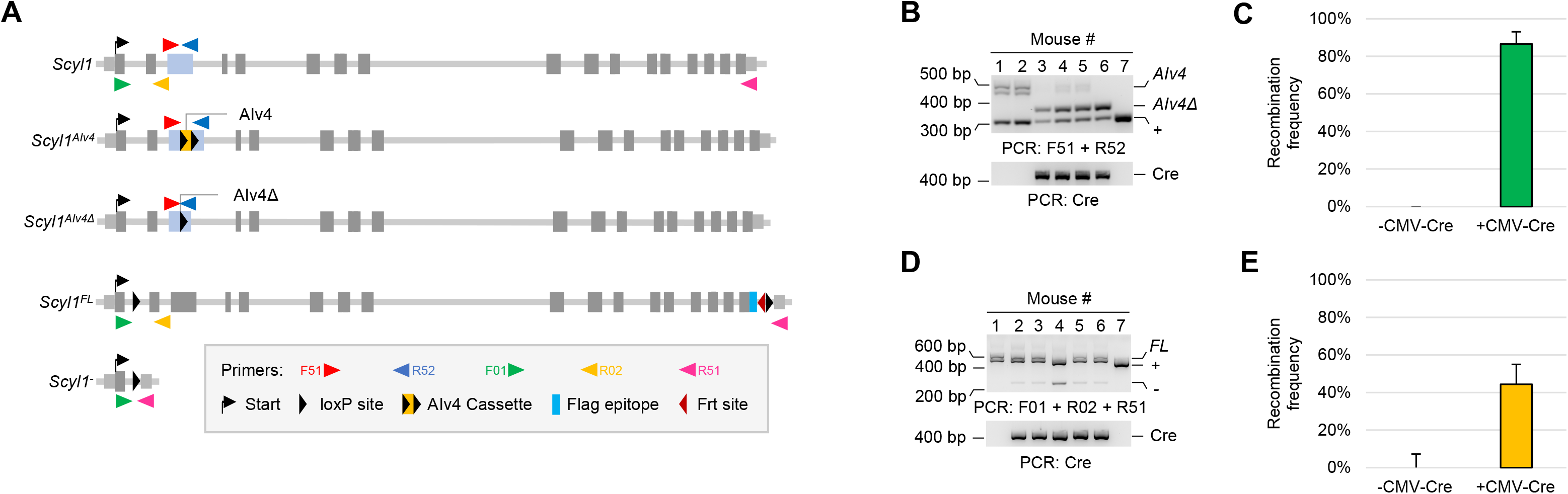
Efficient recombination between the two loxP sites within the AIv4 cassette in vivo. A) Schematic representation of *Scyl1*^+^, *Scyl1^AIv4^*, *Scyl1^AIv4Δ^*, *Scyl1^FL^*, and *Scyl1*^-^ alleles. Dark gray boxes represent exons, with exon 3 highlighted in blue. Light gray boxes represent the 5’ and 3’ untranslated regions of the gene. Red, blue, green, yellow, and pink triangles represent genotyping primers F51, R52, F01, R02, and R51 respectively. The black arrowheads represent the start sites. The yellow box flanked by two black triangles (loxP sites) represents the artificial intron inserted into exon 3 of Scyl1. The light blue box represents a Flag epitope. The crimson triangle represents a Frt site. B) PCR based genotyping of mice obtained from CMV-Cre+ mice crossed with *Scyl1^AIv4/AIv4^* mice. Amplicons of 304, 358, and 505 bp correspond to the wild-type, AIv4, and AIv4Δ alleles, respectively. C) Quantification of recombination of the AIv4 allele into the AIv4Δ allele. Data is expressed as mean ± SEM. D) PCR based genotyping of CMV-Cre+ mice crossed with *Scyl1^FL/FL^* mice. Amplicons of 521,251, and 625 bp corresponding to the wild-type (*Scyl1*^+^), null (*Scyl1*^-^), and floxed alleles (*Scyl1^FL^*), respectively, are obtained. E) Quantification of recombination of the floxed allele into the null allele. Data is expressed as mean ± SEM. Note the increased recombination efficiency in *Scyl1*^+/*AIv4*^ mice compared to *Scyl1*^+/*FL*^ mice.

To test whether recombination between the two loxP sites would result in the inactivation of the *Scyl1* gene, we crossed *CMV-Cre+;Scyl1^+/AIv4Δ^* mice with *Scyl1^AIv4/AIv4^* mice to generate CMV-*Cre+;Scyl1^AIv4Δ/AIv4Δ^* mice (Fig. 4A). From this cross, 36 mice were obtained, 16 of which expressed Cre and 10 were also homozygous for the AIv4Δ allele (62.5% of Cre expressing mice) (Fig. 4B and 4C). PCR fragments corresponding to the AIv4 and AIv4Δ alleles were analyzed by Sanger sequencing and confirmed the proper rearrangement of the recombined allele in Cre expressing mice (Fig. 4D).

**Figure 4.**
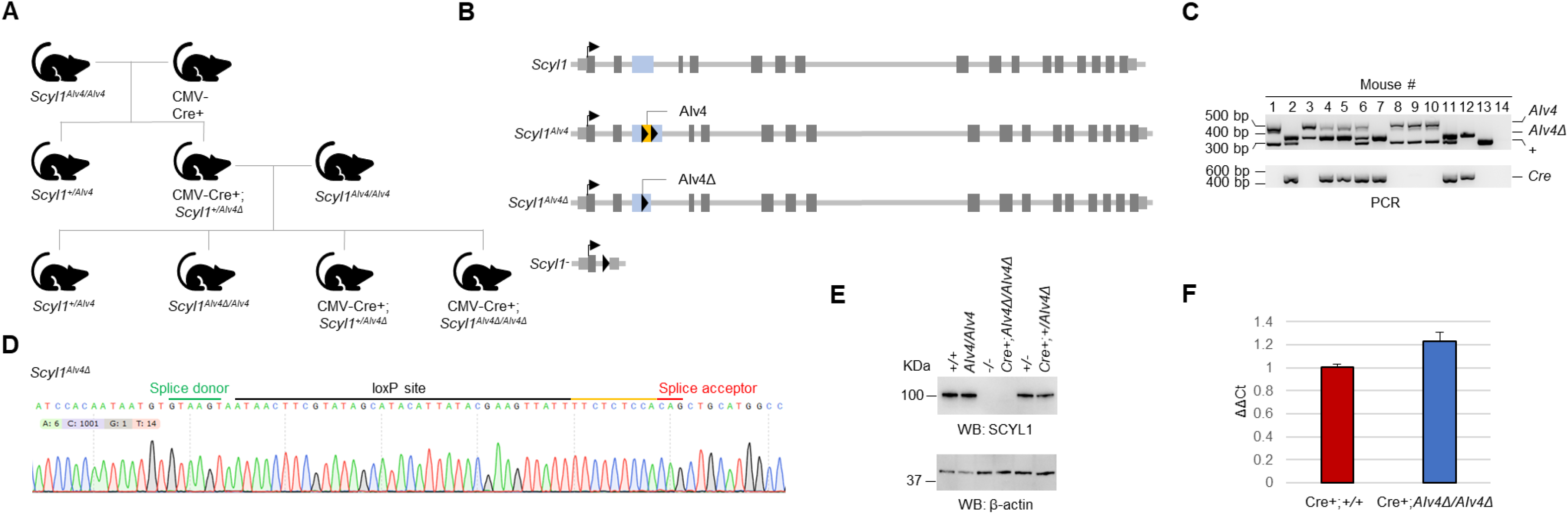
Cre-mediated disabling of the artificial intron leads to complete inactivation of *Scyl1*. A) Breeding scheme for the generation of CMV-Cre+;*Scyl1^AIv4Δ/AIv4Δ^* mice. B) Schematic representation of *Scyl1* alleles generated from this cross: *Scyl1^+^, Scyl1^AIv4^*, and *Scyl1^AIv4Δ^*. For comparison, *Scyl1*^-^ is also illustrated. C) PCR-based genotyping of *Scyl1^+^, Scyl1^AIv4^*, and *Scyl1^AIv4Δ^* alleles. Amplicons of 304, 358, and 505 bp correspond to the wild-type, AIv4, and AIv4Δ alleles, respectively. PCR primers for genotyping are illustrated in Fig. 3A. D) Sanger sequencing was used to confirm proper recombination between the two loxP sites. E) SCYL1 expression in the brain of *Scyl1^+/+^, Scyl1^AIv4/AIv4^, Scyl1^-/-^*, CMV-Cre+;*Scyl1^AIv4Δ/AIv4Δ^, Scyl1^+/-^*, and CMV-Cre+;*Scyl1A*^+/*AIv4Δ*^ mice, as detected by western blotting (top panel), using the anti-SCYL1 7645-AP antibody. β-actin western blot (bottom panel) serves as loading control. F) *Scyl1* gene expression in the cerebella of CMV-Cre+;*Scyl1*^+/+^ and CMV-Cre+;*Scyl1^AIv4Δ/AIv4Δ^* mice. Data is expressed as mean ± SEM.

Importantly, CMV-Cre+;*Scyl1^AIv4Δ/AIv4Δ^* mice, like *Scyl1^-/-^* mice, did not express SCYL1, as revealed by western blotting using an antibody against the carboxyl-terminal segment of SCYL1 (7645-AP) (Fig. 4E). Similarly, no expression of the full-length protein or shorter aberrant isoforms were observed in CMV-*Cre+;Scyl1^AIv4Δ/AIv4Δ^* mice when using an antibody against the amino-terminus of SCYL1 (not shown). These results indicated that crippling the artificial intron via Cre-mediated recombination efficiently inactivated the allele and abrogated SCYL1 expression. Intriguingly, *Scyl1* expression at the mRNA level appeared to be unaffected by the inactivation of the artificial intron (Fig. 4F), suggesting that abrogation of SCYL1 protein expression may be the result of early translation termination rather than degradation of the mRNA transcript by the nonsense-mediated mRNA decay pathway.

Previous studies in our laboratory have shown that inactivation of *Scyl1* results in growth retardation and severe motor dysfunction due to the progressive loss of spinal motor neurons (Kuliyev et al., 2018; Pelletier et al., 2012). To test whether crippling the artificial intron would result in a loss-of-function phenotype, we monitored the weight, grip strength, and visually assessed motor dysfunction using a previously established scoring system (Kuliyev *et al*., 2018; Pelletier *et al*., 2012). Histology was also performed in the hindlimbs of these animals to look for signs of neurogenic atrophy. Consistent with complete inactivation of *Scyl1*, CMV-Cre+;*Scyl1^AIv4Δ/AIv4Δ^* mice exhibited growth retardation (Fig 5A and B), muscle weakness (Fig. 5C) and muscle wasting (Fig. 5C and 5E), as well as sensory motor deficits (Fig. 5D). Muscle fibers in CMV-Cre+;*Scyl1^AIv4Δ/AIv4Δ^* mice showed evidence of neurogenic atrophy such as small angulated fibers, centrally localized nuclei, and group atrophy, similar to *Scyl1-*deficient mice (Fig. 5F) (Kuliyev *et al*., 2018; Pelletier *et al*., 2012).

**Figure 5.**
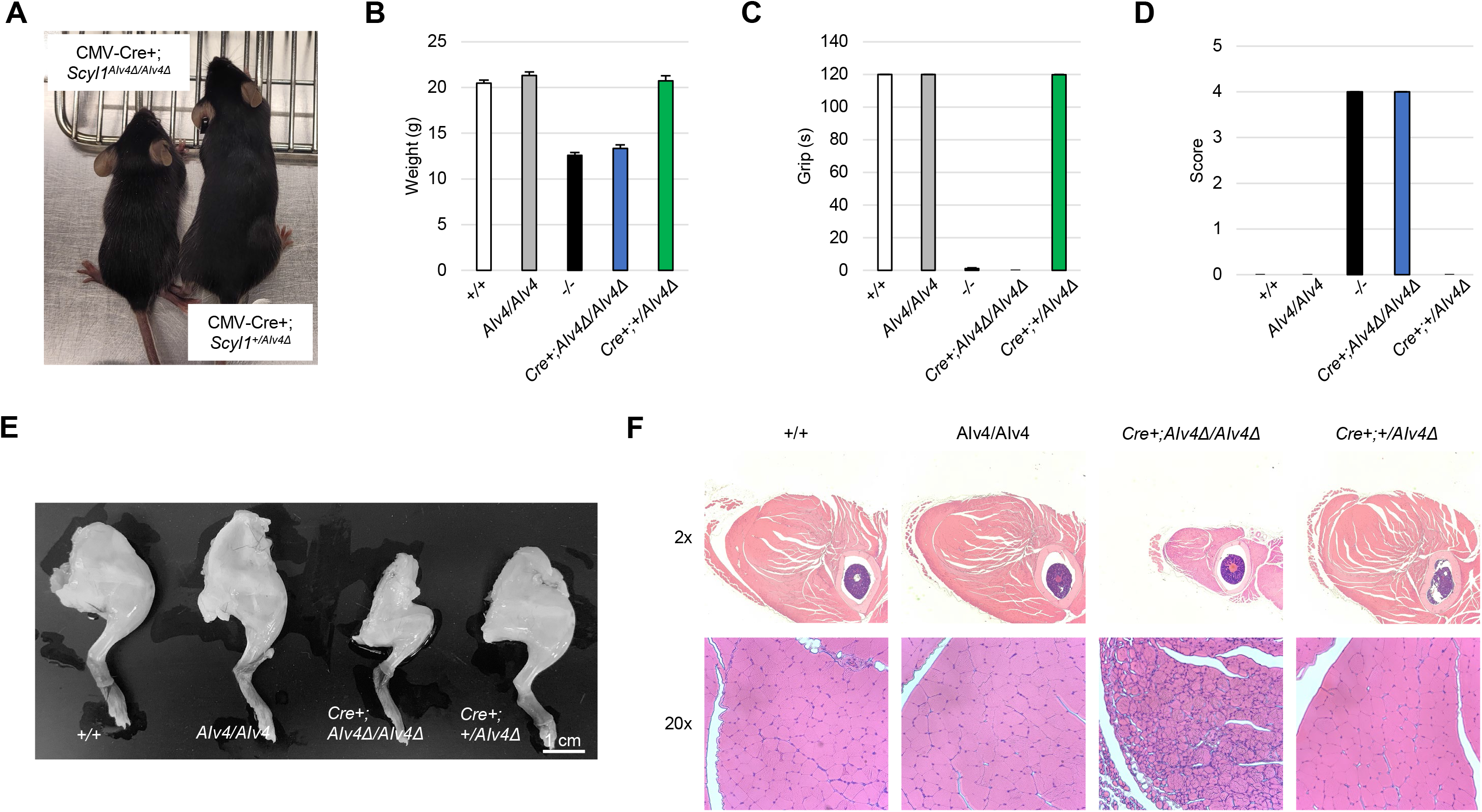
Complete inactivation of the AIv4 allele results in *Scyl1* loss-of-function phenotypes. A) Representative photograph of CMV-Cre+;*Scyl1^AIv4Δ/AIv4Δ^* and CMV-Cre+;*Scyl1^+/AIv4Δ^* mice. Note the size difference between the littermates. B-D) Loss-of-function phenotypes in CMV-Cre+;*Scyl1^AIv4Δ/AIv4Δ^* mice. Similar to *Scyl1*-deficient mice, CMV-Cre+; *Scyl1^AIv4Δ/AIv4Δ^* mice exhibit growth retardation, sensory motor deficit, and muscle weakness. Data is expressed as mean ± SEM. E) Representative photograph of the right hindlimbs of *Scyl1^+/+^, Scyl1^AIv4/AIv4^, CMV-Cre+;Scyl1^AIv4Δ/AIv4Δ^*, and CMV-Cre+;*Scyl1^+/AIv4Δ^* mice. F) Representative micrographs of H&E-stained sections of rectus femoris from 8-week-old *Scyl1^+/+^, Scyl1^AIv4/AIv4^*, CMV-Cre+;*Scyl1^AIv4Δ/AIv4Δ^*, and CMV-Cre+;*Scyl1^+/AIv4Δ^* mice. Note the presence of angulated/atrophied fibers in CMV-Cre+;*Scyl1^AIv4Δ/AIv4Δ^* mice.

The presence of a splice donor site in the AIv4 cassette prompted us to test whether aberrant splicing events with downstream exons or with cryptic splice acceptor sites occurred. To this end, a pair of primers was designed to amplify sequences spanning exons 2 to 5 of the *Scyl1* gene. Intriguingly, RT-PCR reactions performed on total RNA samples from CMV-Cre+;*Scyl1^+/AIv4Δ^* and CMV-Cre+;*Scyl1^AIv4Δ/AIv4Δ^* revealed the presence of shorter-than-predicted PCR fragments that corresponded to alternative splicing events (Fig. 6A). These shorter PCR fragments were not observed in *Scyl1*^+/+^, *Scyl1^+/AIV4^, Scyl1^AIv4/AIv4^* or CMV-Cre+;*Scyl1*^+/+^ mice, indicating that they likely resulted from the Cre-mediated recombination of the AIv4 intron. TOPO cloning and Sanger sequencing of the RT-PCR fragments revealed the presence of multiple splicing events, two of which accounted for 87.5% of the observed transcripts (Fig. 6B). The first involved the splicing between the splice donor of AIv4 and the splice acceptor of exon 4 (transcript variant 1); the second splicing event occurred between the AIv4 splice donor and a cryptic splice acceptor within the 3’ half of exon 3 (transcript variant 2). These splicing events, interestingly, are in-frame suggesting that a shorter, mutant version of SCYL1 may still be expressed. Western blot analyses, however, revealed the complete absence of SCYL1 in the brain of CMV-Cre+;*Scyl1^AIv4Δ/AIv4Δ^* animals, suggesting that these mutant protein isoforms may be unstable and thus rapidly degraded after synthesis. Acknowledging the existence of such in-frame splicing events is essential for proper design of additional alleles using this approach. Design considerations are presented in the discussion.

**Figure 6.**
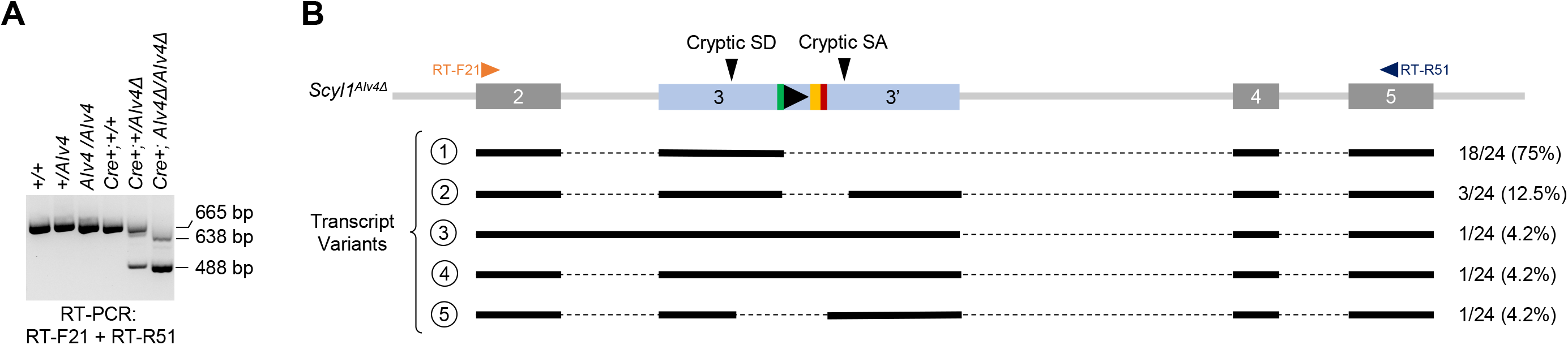
Transcript variants produced from the AIv4Δ allele. A) RT-PCR products generated from cerebellar total RNA extracts of *Scyl1^+/+^, Scyl1^+/AIv4^, Scyl1^AIv4/AIv4^*,CMV-Cre+;*Scyl1*^+/+^, CMV-Cre+;*Scyl1^+/AIv4Δ^*, and CMV-Cre+;*Scyl1^AIv4Δ/AIv4Δ^* mice. Amplicons of 665, 638, and 488 bp, corresponding to the mature form of the *Scyl1* transcript, as well as shorter transcript variants 1 and 2 were obtained. These shorter transcripts were observed only in CMV-Cre+;*Scyl1^+/AIv4Δ^* and CMV-Cre+;*Scyl1^AIv4Δ/AIv4Δ^* mice. B) Schematic representation of the five most abundant RNA transcripts observed in CMV-Cre+;*Scyl1^AIv4Δ/AIv4Δ^* mice (transcripts variants labeled 1-5). RT-PCR products from CMV-Cre+;*Scyl1^AIv4Δ/AIv4Δ^* mice were TOPO cloned. Plasmid DNA was recovered from 24 clones and analyzed by Sanger sequencing. The frequency of each transcript is illustrated to the right of the transcript. 18 out of 24 clones were obtained for transcript variant 1; 3 out of 24 clones were obtained for transcript variant 2; and 1 out of 24 clones were obtained for transcript variants 3, 4, and 5. Transcript variant 1 resulted from the splicing of the AIv4 splice donor site into the splice acceptor site of exon 4. Transcript variant 2 resulted from the splicing of the AIv4 splice donor site into a cryptic splice acceptor site within the 3’ half of exon 3. Transcript variant 3 contained intronic sequences from intron 2 and the recombined AIv4 cassette as well as sequences encoding exon 2 to 5. Transcript variant 4 contained the expected sequence of the recombined AIv4 cassette. Transcript variant 5 resulted from the splicing of a cryptic splice donor site within the 5’ half of exon 3 into the AIv4 splice acceptor site.

## Discussion

Here we describe the successful application of the DECAI approach for the generation of conditional alleles in mice via zygote injection of CRISPR-Cas9 components. The approach is straightforward, effective, and results in complete inactivation of the allele upon Cre-mediated recombination. The approach also provides several advantages over commonly used strategies and may represent a faster and more affordable solution to engineer conditional alleles in mice. Some modifications to the original strategy were also introduced to simplify the engineering process and improve efficiency. Finally, considering our findings, we provide guidelines for the design of alleles using this approach.

Currently, there are two main strategies employed to engineer conditional alleles in mice: the flanking of one or more critical exons with SSRSs or the insertion of large modules within an exon of a gene. Both design strategies can be used to engineer conditional alleles in mice via conventional or CRISPR-assisted gene targeting in ES cells or via microinjection of CRISPR components directly into pronuclear stage zygotes. While these approaches have been used to engineer several thousands of mouse models over the past decades, these approaches are labor intensive and inefficient. Flanking critical exons using conventional or CRISPR-assisted gene targeting in ES cells involves the generation of large targeting constructs and often requires screening of several hundred clones. These approaches also often involve selection markers, leaving genetic scars at the engineered loci. Moreover, several rounds blastocyst injections are often needed to obtain chimeric animals capable of transmitting the genetic alteration. Flanking exons with SSRSs via zygote microinjection of CRISPR reagents and short ssDNA molecules encoding SSRSs flanked by short homology arms also has its share of limitations. SSRSs must be inserted in cis which significantly reduces the chance of generating floxed alleles (Pelletier *et al*., 2015). Although the use of large DNA molecules encoding both SSRSs were shown to improve this process (Codner et al., 2018; Miura et al., 2018; Quadros et al., 2017), their synthesis is expensive, frequently unsuccessful, and the molecules are often too small to flank multiple exons. Additionally, the approach of flanking exons with SSRSs cannot be used to inactivate single exon or overlapping genes, as the insertion of SSRSs may disrupt regulatory elements within the promoter or the 3’ UTR of those genes. The use of large artificial introns for engineering conditional alleles in mice also share many of the aforementioned constraints, as these are typically used for gene targeting in ES cells and require the generation of large, complex targeting constructs (Andersson-Rolf *et al*., 2017; Economides *et al*., 2013). Moreover, these rely on the activity of a splice acceptor site which can be leaky, allowing for expression of the target gene, as well as strong promoters within reporter cassettes, which may affect the expression of surrounding genes (Andersson-Rolf *et al*., 2017; Economides *et al*., 2013; Soulez *et al*., 2019).

The modified DECAI approach described here makes use of a short 319-nucleotide-long ssDNA molecule as an HDR template to insert a 201-nucleotide-long cassette. Short ssDNA molecules are inexpensive, generally easy to synthesize, and are efficiently inserted via microinjection of CRIPSR reagents into pronuclear stage zygotes (Pelletier *et al*., 2015), bypassing the need to engineer ES cells. Insertion of the small artificial intron used in our study occurred in 16.7% of edited pups obtained from zygote injection. More recently, we obtained an insertional rate of 50% for another gene, indicating that a single round of injection may be sufficient to generate conditional alleles in mice as opposed to 2 to 4 rounds of injections required for current methodologies (Pelletier *et al*., 2015). The genotyping of such an allele is also greatly simplified compared to genotyping floxed alleles or conditional alleles generated using large molecules as it only requires one pair of primers, PCR amplification of the locus, and Sanger sequencing. Another foreseeable advantage of using this technology to engineer conditional alleles in mice is that, like most CRISPR-based approaches, it can be multiplexed and several mouse models with conditional alleles can be engineered from a single round of microinjection. Alleles can be subsequently segregated by outbreeding founder mice with wild type mice. This approach is currently being tested in our facility. If successful, multiplexing may again significantly reduce the cost and time required for engineering conditional alleles in mice. Although not tested here, the approach can also be used to conditionally inactivate single exon genes as well as overlapping genes as the modification affects gene expression at the mRNA and/or translational level rather than at the structural level.

Despite the numerous advantages of this approach, some potential limitations to the use of this technology may exist. For example, the preference of GT-AG introns for specific sequences upstream and downstream of the splice donor and acceptor sites may limit the number of possible target sites for insertion of the intron (Sibley et al., 2016). These sequences generally contain a CAG or AAG upstream of the donor site and an A or G immediately downstream of the splice acceptor site. However, when engineering the *Scyl1^AIv4^* allele, this notion was overlooked and the artificial intron was inserted two nucleotides upstream of the SpCas9 cut site, immediately after a TGT and before a C. While these sequences are thought to be suboptimal to promote splicing (Sibley *et al*., 2016), SCYL1 expression in *Scyl1^AIv4/AIv4^* and *Scyl1*^+/+^ mice was similar, indicating that the artificial intron is efficiently removed by the splicing machinery. Thus, limiting the insertion of the AIv4 cassette to CAG-A/G or AAG-A/G sites may not be required for engineering conditional alleles using this approach.

Another potential downside of using this technology is that insertion of the intron may affect gene expression by deregulating endogenous splicing events (Guzzardo *et al*., 2017). Although RT-qPCR and western blotting results presented here indicate that no such dysregulation occurred in *Scyl1^AIv4/AIv4^* mice, aberrant splicing events in CMV-Cre+;*Scyl1^+/AIv4Δ^* and CMV-Cre+;*Scyl1^AIv4Δ/AIv4Δ^* mice were observed. These aberrant splicing events were only observed when the AIv4 polypyrimidine tract and the branch point were removed via Cre-mediated recombination, indicating that crippling of the intron leaves behind an active splice donor site that drives unpredictable and potentially undesirable splicing events. This observation is extremely important and should not be disregarded especially if these splicing events are in-frame, as a mutant form of the protein with aberrant properties may be expressed. Fortunately, no mutant forms of SCYL1 were detected by western blotting using antibodies against the amino-terminal or the carboxyl-terminal segment of SCYL1, suggesting that if truncated forms of SCYL1 were produced, they were likely unstable and immediately degraded after synthesis. Consistent with the generation of a loss-of-function allele, CMV-Cre+;*Scyl1^AIv4Δ/AIv4Δ^* also exhibited growth retardation and motor dysfunction.

Two main splicing events were identified in CMV-Cre+;*Scyl1^AIv4Δ/AIv4Δ^* mice. The first was a splicing event between the splice donor of AIv4 and the splice acceptor of exon 4 (transcript variant 1), which accounted for 75% of the transcripts. The second was the splicing between the AIv4 splice donor and a cryptic splice acceptor site located within the 3’ half of exon 3 (transcript variant 2), which accounted for 12.5% of all transcripts. Both splicing events resulted in in-frame and thus functional transcripts which could have resulted in the expression of SCYL1 isoforms with aberrant properties. Although this was not the case, we suspect that the vast majority of the RNA transcript detected by RT-qPCR were either transcript variant 1 or 2. The low representation of the out-of-frame variants, including the expected transcript containing the recombined AIv4 allele (transcript variant 4), may result from their degradation via the nonsense-mediated mRNA decay pathway. In light of these findings, we propose the following guidelines for the generation of conditional alleles using AIv4:

1. The artificial intron should be inserted within the 5’ most exons of a gene to promote mRNA degradation via the nonsense-mediated mRNA decay pathway (Popp and Maquat, 2016).
2. The artificial intron should be inserted within the 5’ half of an exon to promote mRNA degradation via the nonsense-mediated mRNA decay pathway (Popp and Maquat, 2016).
3. The artificial intron should be inserted such that potential splicing events driven by the AIv4 splice donor with downstream exon(s) results in an out of frame transcript.
4. The artificial intron can be inserted anywhere, regardless of optimal sequences upstream and downstream of the splice donor and acceptor sites.
5. Short homology arms (of approximately 60 nucleotides) can be used to promote insertion.
6. ssDNA molecules should be used for the generation of mouse models as it is less toxic than double stranded molecules and does not randomly integrate into the genome.

In conclusion, we believe that the application of this technology may significantly reduce the cost and time required for making mouse models with conditional alleles. Design considerations such as those presented herein are essential to the successful application of this technology.

## Acknowledgements

The authors would like to thank members of the Indiana University Genome Editing Center (IUGEC) for rederiving the *Scyl1^AIv4^* mouse model, the St. Jude children’s Research Hospital Transgenic and knockout core facility for microinjections, and the Histology Core of the Indiana Center for Musculoskeletal Health at IU School of Medicine and the Bone and Body Composition Core of the Indiana Clinical Translational Sciences Institute (CTSI) for skeletal muscle histology. Funding supporting this work was provided by the Indiana University School of Medicine and the Department of Medical and Molecular Genetics.

## Methods

### Generation of Scl1^AIv4^ mice

*Scyl1^AIv4^* mice were engineered using CRISPR-Cas9 technology. Briefly, C57BL/6J zygotes were injected with Cas9 mRNA transcript (100 ng/μL), a sgRNA targeting exon 3 of *Scyl1* (50 ng/μL), and an HDR template in the form of a ssDNA molecule (1.2 pmol/μL, see Table 1 for complete sequences). Zygotes were then transferred to pseudopregnant CD1 females. Pups obtained from these microinjections were characterized by PCR-based genotyping and Sanger sequencing using the following primers: S1E154D-F1 5’-TACTCTCCTCAGGCCCTCAC-3’ and S1E154D-R1 5’-GAAGCACCGAACACCCAAAC-3’. A fragment of 667 bp was obtained for the wild-type allele, while the AIv4 allele produced a fragment of 868 bp. Sanger sequencing was performed using both forward and reverse primers. Guide sequence selection, HDR template design, and synthesis of Cas9 mRNA and sgRNA transcripts were performed as previously described in (Cassidy et al., 2022). The following primer was used for synthesis of the sgRNA: S1-HAN-Guide_01-F 5’-TAATACGACTCACTATAGGCATCCACAATAATGTCTGCAGTTTTAGAGCTAGAAATAGCA-3’

### Sanger sequencing

Sanger sequencing was performed as previously described in (Cassidy *et al*., 2022).

### Previously described mouse models used in this study

B6.C-Tg(CMV-cre)1Cgn/J, 006054 mice (CMV-Cre+) were purchased from Jackson Laboratory (Bar Harbor, ME). *Scyl1^FL^* mice were generated in our laboratory and were previously described in (Pelletier *et al*., 2012).

### Generation of CMV-Cre+;Scyl1^Alv4Δ^ mice

CMV-Cre+;*Scyl1^AIv4Δ/AIv4Δ^* mice were generated first by crossing *Scyl1^AIv4/AIv4^* females with CMV-Cre+ males (B6.C-Tg(CMV-cre)1Cgn/J, 006054). From this cross, we obtained CMV-Cre+;*Scyl1^+/AIv4Δ^* females, which were then bred with *Scyl1^AIv4/AIv4^* males.

### PCR Genotyping

CMV-Cre+ mice were routinely genotyped using the following primers: Cre66 5’-CCTGCGGTGCTAACCAGCGTT-3’ and Cre99 5’-TGGGCGGCATGGTGCAAGTT-3’. Mice positive for Cre recombinase produced a fragment of 470 bp, whereas no fragments were obtained from wild-type mice. Mice bearing *Scyl1^AIv4^* and *Scyl1^AIv4Δ^* alleles were routinely genotyped using the following primers: Scyl1_AIv4_F51 5’-GTGCCTCCACATCGTGACAG-3’ and Scyl1_AIv4_R52 5’-CTCCGGGGGATCATACTGCT-3’. Fragments of 505, 358, and 304 bp corresponding to the *Scyl1^AIv4^, Scyl1^AIv4Δ^*, and wild-type alleles were obtained. Mice bearing the *Scyl1^FL^* and *Scyl1*^-^ were routinely genotyped using the following primers: S1F01 (5’-GCTGCTCCGAAGGCCGCGGCCGA-3’), S1R51 (5-GATTATGTACACTAGATGTGCCTGA-3’), and S1R02 (5’-GAGGAGAGTAAGATGGGTAGA-3’). Bands of 521, 251, and 625 bp corresponding to the wild-type (*Scyl1*^+^), null (*Scyl1*^-^), and floxed alleles (*Scyl1^FL^*), respectively, were obtained. All PCR genotyping reactions were performed using Taq DNA polymerase from Qiagen (201205).

### Western blot

SCYL1 expression in the brains of *Scyl1*^+/+^, *Scyl1*^+/-^, *Scyl1*^-/-^, *Scyl1^AIv4/AIv4^*, *CMV-Cre^+^;Scyl1^AIv4Δ/AIv4Δ^*, and CMV-Cre+;*Scyl1^+/AIv4Δ^* was analyzed by western blot. Briefly, brains were harvested, rinsed with PBS and snap frozen on dry ice. The brains were then lysed in Triton X-100 buffer [50 mM Tris-HCl, pH 8.0, 150 mM NaCl, 1% TX-100, PhosSTOP™ Phosphatase Inhibitor Cocktail tablets (4906837001, Sigma-Aldrich) and cOmplete™, EDTA-free Protease Inhibitor Cocktail tablets (4693132001, Sigma-Aldrich)]. Samples were cleared by centrifugation at 4°C for 30 minutes, 5,000 x g and proteins were quantified using the Peirce BCA Assay (23225, Life Technologies). 5 μg of total protein were loaded on 10% Criterion™ XT Bis-Tris Protein Gels for separation. The resolved proteins were then transferred to LF-PVDF membranes and probed with an anti-SCYL1 antibody or b-actin antibody overnight at 4°C in TBST (137 mM sodium chloride, 20 mM Tris, 0.1% Tween 20, pH 7.6) supplemented with 3% nonfat dry milk. The next day, the membranes were washed three times and probed with secondary antibodies for 1 hour at room temperature. Following incubation with the secondary antibodies, the membranes were washed three times and the proteins detected by chemiluminescence or fluorescence using the ChemiDoc™ MP Imaging System.

### Animal Phenotyping

Phenotypic abnormalities in *Scyl1*^+/+^, *Scyl1^AIv4/AIv4^*, *Scyl1*^-/-^, CMV-Cre+;*Scyl1^AIv4Δ/AIv4Δ^*, and CMV-Cre+;*Scyl1^+/AIv4Δ^* mice were evaluated at 8 weeks of age. Growth abnormalities were measured by weighing the mice. Muscle strength was monitored by using an inverted cage grid test, as described previously (Pelletier *et al*., 2012). Additionally, motor dysfunction was assessed using a previously established scoring system where a score of 1 indicates visible growth abnormalities; 2 indicates growth defects and abnormal gait; 3 indicates posterior waddle and abnormal gait; 4 indicates growth abnormalities, abnormal gait, and tremor when suspended by their tails; 5 indicates aforementioned phenotypes and partial paralysis; 6 indicates complete paralysis; and 7 indicates flattening of the pelvis and dorsally contracted hindlimbs (Pelletier *et al*., 2012). For the grip test, mice were placed on top of an elevated cage grid. When mice were holding tightly to the cage grid, the grid was inverted and then the amount of time that they remained suspended (maximum 120 seconds) was recorded.

### Histology

Histologic studies were performed on tissues collected from 3 females from each of the following genotypes: *Scyl1*^+/+^, *Scyl1^AIv4/AIv4^*, CMV-Cre+;*Scyl1^AIv4Δ/AIv4Δ^*, and CMV-Cre+;*Scyl1^+/AIv4Δ^*. Immediately after the animals were sacrificed, right hindlimbs were postfixed by immersion in 4% paraformaldehyde in PBS for at least 24 hours at 4°C before being decalcified in formic acid. Tissues were embedded in paraffin, sectioned at 8 μm, mounted on positively charged glass slides, and dried in a 60°C oven for 20 min. Slides were stained with hematoxylin and eosin (H&E) using LINISTAIN GLX linear stainer. Images of H&E-stained limb cross-sections were acquired with an Echo Revolve microscope.

### RT-qPCR

*Scyl1* expression was analyzed at the mRNA level using RT-qPCR. Briefly, cerebella were harvested from CMV-Cre+;*Scyl1^AIv4Δ/AIv4Δ^* and CMV-Cre+;*Scyl1*^+/+^ mice, rinsed with PBS and snap frozen in liquid nitrogen. Total RNA was extracted using Rneasy Maxi Kit (Qiagen, 75162) with the RNase-Free DNase Set (Qiagen, 79254). The quality of the RNA was assessed using the Agilent TapeStation system and quantified by Nanodrop. cDNA was produced from 100 ng of total RNA using the ProtoScript^®^ II First Strand cDNA Synthesis Kit (New England Biolabs, E6560L) with the d(T)23 VN primer, according to the manufacturer’s recommendations. *Scyl1* expression was quantified using TaqMan 2X Master Mix (Applied Biosystems, 4444557) and TaqMan Gene Expression Assay Probe Mm00452459_m1(FAM), which was normalized against *Gapdh* expression using TaqMan Gene Expression Assay Probe Mm99999915_g1 (FAM). Relative expression of *Scyl1* was determined using the ΔΔCt method.

### RT-PCR

The detection of aberrant *Scyl1* transcripts was performed by RT-PCR using primers designed to amplify sequences between exon 2 to exon 5 (RT-Scyl1_F21 5’-CGCAGTGTCCATCTTCGTGTA-3’ and RT-Scyl1_R51 5’-CCCGGCAGTTCTGCAGGAA-3’) following the OneTaq One-Step RT-PCR Kit (New England Biolabs, E5315S) procedure. Briefly, cerebella were harvested from *Scyl1*^+/+^, *Scyl1*^+/*AIv4*^, *Scyl1^AIv4/AIv4^*, CMV-Cre+;*Scyl1*^+/+^, CMV-Cre+;*Scyl1*^+/*AIv4*^, and CMV-Cre+;*Scyl1^AIv4Δ/AIv4Δ^* mice, rinsed with PBS and snap frozen in liquid nitrogen. Total RNA was extracted using Rneasy Maxi Kit (Qiagen, 75162) and the additional RNase-Free DNase digestion step (Qiagen, 79254). The quality and quantity of the RNA were assessed as described above.

### Mouse Husbandry

Mice were housed in an Association for Assessment and Accreditation of Laboratory Animal Care accredited facility and maintained in accordance with the National Institutes of Health Guide for the Care and Use of Laboratory Animals. Animal experiments were reviewed and approved by the Indiana University Institutional Animal Care and Use Committee.

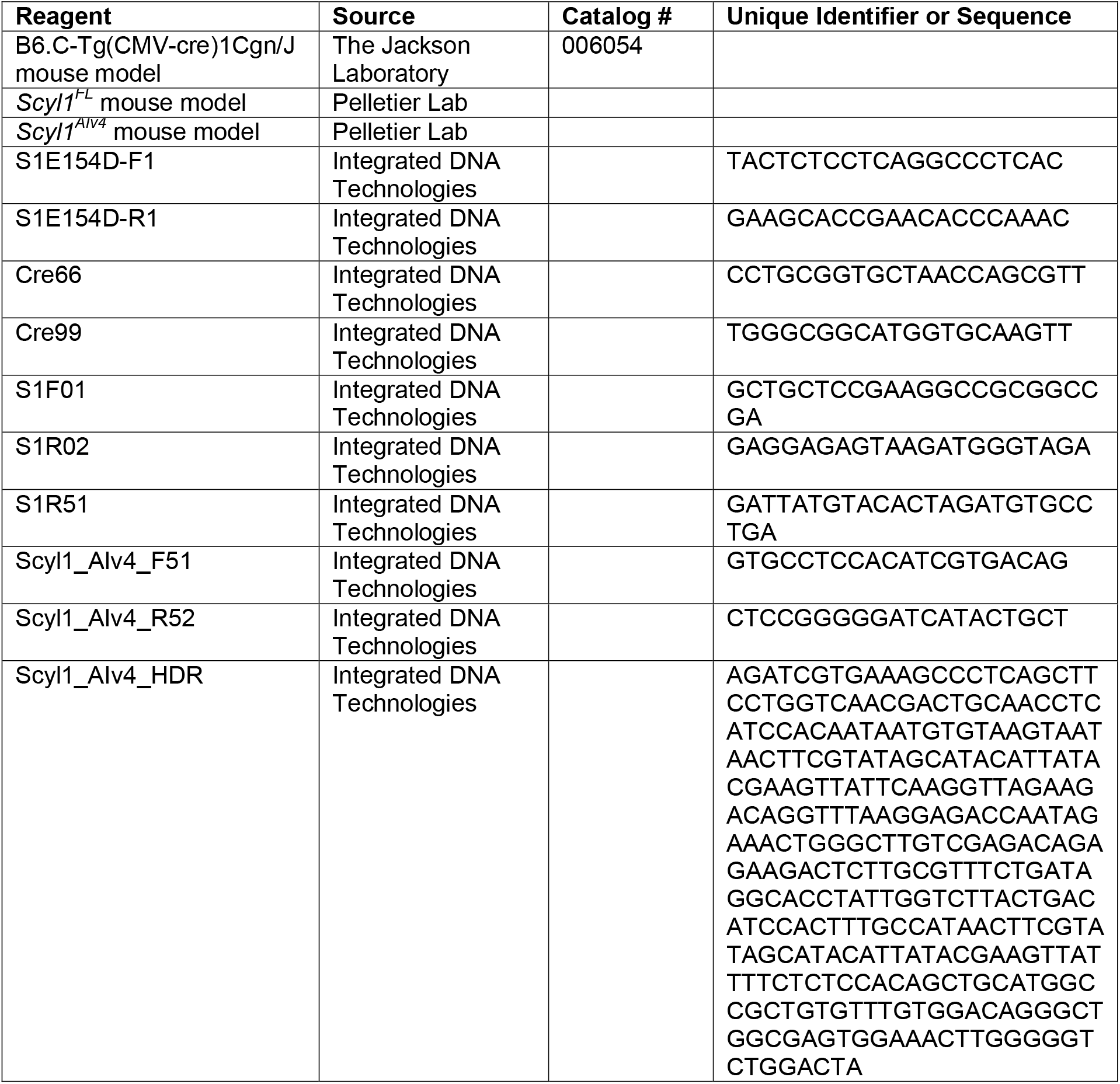
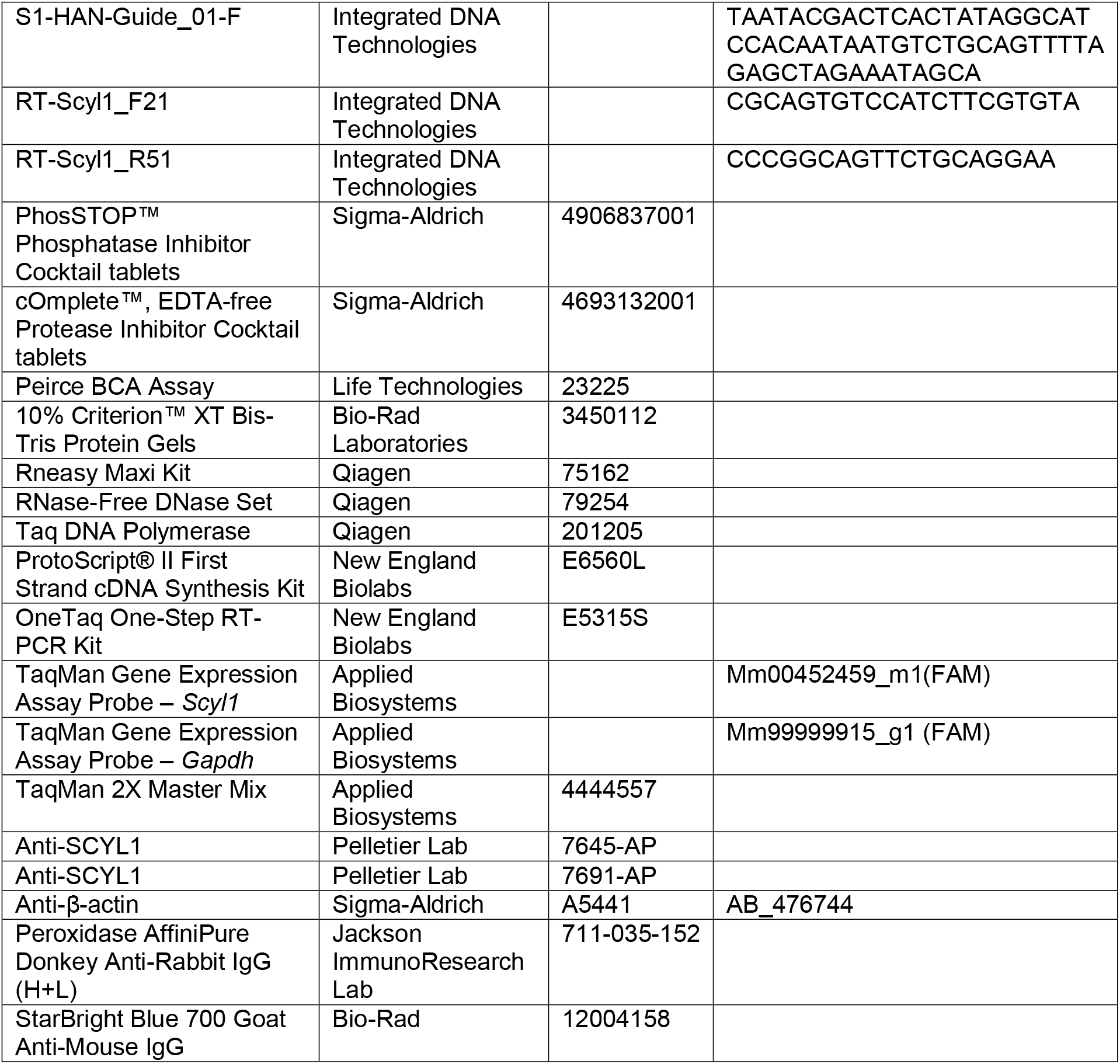

